# Kaposi’s sarcoma-associated herpesvirus ORF68 is a DNA binding protein required for viral genome cleavage and packaging

**DOI:** 10.1101/322719

**Authors:** Matthew R Gardner, Britt A Glaunsinger

## Abstract

Herpesviral DNA packaging into nascent capsids requires multiple conserved viral proteins that coordinate genome encapsidation. Here, we investigated the role of the ORF68 protein of Kaposi’s sarcoma-associated herpesvirus (KSHV), a protein required for viral DNA encapsidation whose function remains largely unresolved across the herpesviridae. We found that KSHV ORF68 is expressed with early kinetics and localizes predominantly to viral replication compartments, although it is dispensable for viral DNA replication and gene expression. However, in agreement with its proposed role in viral DNA packaging, KSHV-infected cells lacking ORF68 failed to cleave viral DNA concatemers, accumulated exclusively immature B-capsids, and released no infectious progeny virions. ORF68 has no predicted domains aside from a series of putative zinc finger motifs. However, *in vitro* biochemical analyses of purified ORF68 protein revealed that it robustly binds DNA and is associated with nuclease activity. These activities provide new insights into the role of KSHV ORF68 in viral genome encapsidation.

**Importance:** Kaposi’s sarcoma-associated herpesvirus (KSHV) is the etiologic agent of Kaposi’s sarcoma and several B-cell cancers, causing significant morbidity and mortality in immunocompromised individuals. A critical step in the production of infectious viral progeny is the packaging of the newly replicated viral DNA genome into the capsid, which involves coordination between at least seven herpesviral proteins. While the majority of these packaging factors have been well studied in related herpesviruses, the role of the KSHV ORF68 protein and its homologs remains unresolved. Here, using a KSHV mutant lacking ORF68, we confirm its requirement for viral DNA processing and packaging in infected cells. Furthermore, we show that the purified ORF68 protein directly binds DNA and is associated with a metal-dependent cleavage activity on double stranded DNA *in vitro*. These activities suggest a novel role for ORF68 in herpesviral genome processing and encapsidation.

## Introduction

Herpesviruses are large, double-stranded DNA viruses with complex virion assembly and maturation strategies. Following the expression of structural genes, capsids are assembled in the nucleus, first through the production of fragile procapsids (1). Procapsids are formed by the binding and orientation of capsid proteins onto a protein scaffold, which is subsequently digested by the viral protease, allowing further maturation into A-, B-, or C-capsids. A-capsids lack both the protein scaffold and DNA genome and most likely arise from failed packaging events (2). B-capsids retain the cleaved protein scaffold and may represent a DNA-packaging failure, like A-capsids, or simply represent capsids that have not yet encountered the DNA-packaging machinery. Successful packaging of the viral genome, which displaces the digested scaffold proteins, results in the mature, DNA-containing C-capsids (3).

The process of viral DNA replication generates head-to-tail concatemers, which are cleaved into unit-length genomes as they are packaged into immature capsids. In alpha and betaherpesviruses, DNA packaging is coordinated by seven viral proteins, each of which is essential for both cleavage and packaging of the newly replicated genomes (4). The proteins required for DNA packaging have been most extensively studied in the alphaherpesvirus Herpes Simplex virus type 1 (HSV-1), where the essential proteins are UL6, UL15, UL17, UL25, UL28, UL32, and UL33. UL6 forms a ring-shaped oligomer, which occupies one vertex of the icosahedral capsid, forming the portal through which the DNA is threaded during both packaging and unpackaging (5). UL28, UL15, and UL33 form the terminase motor, an ATP-dependent motor that physically threads the viral genome through the portal complex into the nascent capsid (6). Of the terminase motor, only UL28 has been shown to directly bind DNA (7). However, UL15 and its homolog UL89 in betaherpesviruses have been shown to nick DNA substrates *in vitro* (4, 8). UL17 and UL25 are part of the vertex-specific complex, which binds to the vertices of the assembled capsid, as well as to the portal complex, thereby aiding docking of the terminase motor (9, 10). UL32 is the only protein that is required for DNA cleavage and packaging that lacks a well-defined function (4). However, UL32 contains a number of putative C-X-X-C type zinc finger domains and mutation of these conserved motifs results in altered disulfide patterns on the viral protease, which is involved in capsid assembly, supporting a role for UL32 in viral genome encapsidation (11).

Significantly less is known about the DNA packaging process in gammaherpesviruses such as Kaposi’s sarcoma-associated herpesvirus (KSHV), although each of the packaging proteins described above are conserved. These include KSHV ORF7 (UL28; terminase subunit), ORF19 (UL25; vertex-specific complex), ORF29 (UL15; terminase subunit), ORF32 (UL17; vertex-specific complex), ORF43 (UL6; portal protein), ORF67.5 (UL33; terminase subunit), and ORF68 (UL32; unknown). Cryo-electron microscopy of KSHV capsids indicated that KSHV ORF43 forms the portal complex and that ORF32 is associated with vertices, consistent with studies in HSV-1 (12, 13).

Given that role ORF68 and its homologs play during viral DNA packaging are largely unresolved across the herpesviridae, we sought to explore its function in more detail in KSHV-infected cells and *in vitro*. During lytic KSHV infection, many features of ORF68 were conserved with its alpha and betaherpesvirus homologs, such as localization to viral replication compartments, and its requirement for viral genome cleavage, packaging, and infectious virion production. Interestingly, biochemical analyses of recombinant ORF68 protein purified from mammalian cells revealed it to be a DNA-binding protein that is associated with dsDNA cleavage activity. These novel findings suggest that ORF68 plays a previously unappreciated role in KSHV genome processing and encapsidation.

## Results

### KSHV ORF68 is an early viral protein that localizes to viral replication compartments

Although KSHV ORF68 has not been characterized, its homologs in alpha and betaherpesviruses are expressed with late kinetics and are involved in viral DNA packaging (11, 14). ORF68 shares 24.8% sequence identity with UL32, its best characterized homolog from HSV-1, including the five C-X-X-C motifs conserved across all the subfamily homologs (11). To characterize the expression kinetics and potential functions of KSHV ORF68, we generated a rabbit polyclonal antibody using recombinant ORF68 protein. We monitored the expression of ORF68 in KSHV-infected iSLK.BAC16 cells, a commonly used Caki-1 cell line containing a doxycycline (dox)-inducible version of the major viral lytic transactivator, ORF50 (RTA) (15, 16). Lytic reactivation of these cells upon dox treatment revealed that ORF68 expression begins as early as 6 hours post-reactivation and it is robustly expressed by 18 hours (Figure 1a). Its expression kinetics mimicked those of the KSHV delayed early protein ORF59, and it was not impacted by treatment with the viral DNA replication inhibitor phosphonoacetic acid (PAA), which blocks expression of viral late genes such as K8.1 (Figure 1a). Thus, KSHV ORF68 is a delayed early gene.

**Figure 1.**
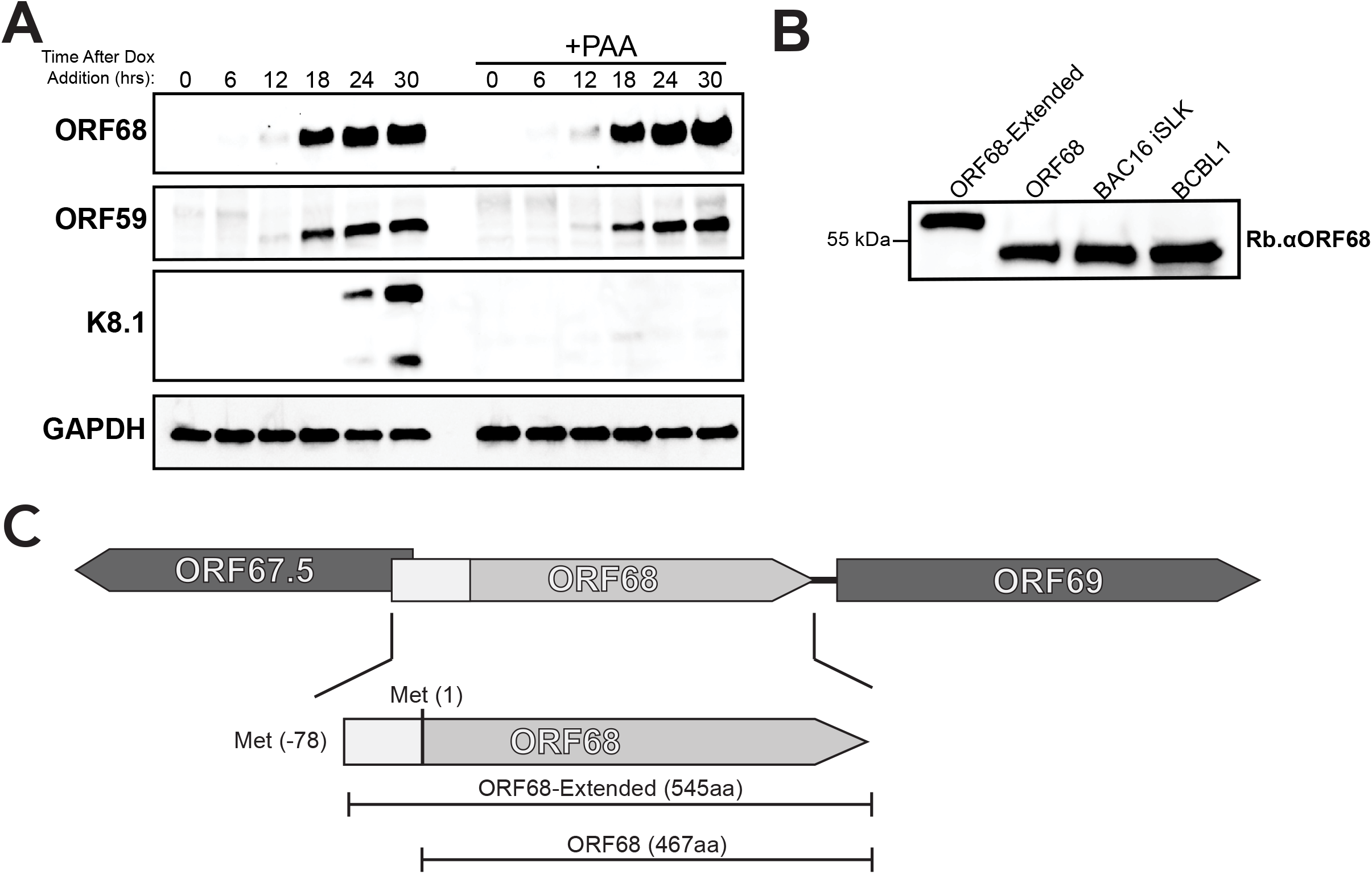
Characterization of the expression of KSHV ORF68. (A) Western blot showing the expression kinetics of ORF68 relative to the early protein ORF59 and the late protein K8.1 in iSLK.BAC16 cells post reactivation with doxycycline (dox). GAPDH serves as a loading control. (B) Anti-ORF68 western blot of lysates of HEK293T cells transfected with plasmids expressing either the extended (ORF68-Extended) or shorter form of ORF68, next to lysates of lytically reactivated iSLK.BAC16 and TREx-BCBL1 cells. Different amounts of lysate were loaded in each lane to account for differing ORF68-expression levels. (C) Diagram showing the genomic locus of ORF68, showing the two annotated forms, which differ by a 78 amino acid (aa) N-terminal extension.

ORF68 has been alternatively annotated as a 467 amino acid (aa) protein and as a longer, 545 aa protein containing a 78 aa N-terminal extension (ORF68-Extended) (Figure 1c). However, on a western blot of iSLK.BAC16 lysate, ORF68 migrated as a single band at the predicted molecular weight (MW) of the 467 aa form (52kDa). To confirm that the 467 aa version was the expressed form in KSHV-infected cells, we cloned both versions into a mammalian expression vector and compared their migration pattern in transfected HEK293T cells to endogenous ORF68 in lytically-reactivated iSLK.BAC16 cells and the KSHV-infected B cell line TREx-BCBL1. Indeed, in both cell lines, only the 467 aa form of ORF68 was detected (Figure 1b).

We next evaluated the subcellular localization of ORF68 in reactivated iSLK.BAC16 and TREx-BCBL1 cells using immunofluorescence assays with the ORF68 antibody. To identify infected cells, cells were co-stained with an antibody against the viral polymerase processivity factor ORF59, which accumulates in viral replication compartments in the nucleus (17). We observed no ORF68 staining in unreactivated or ORF59-negative cells, but in reactivated cells ORF68 was present in both the cytoplasm and, more prominently, in the nucleus. In the cytoplasm, it was primarily diffuse, but also localized to some randomly distributed puncta and larger aggregates close to the nucleus. In the nucleus, ORF68 was localized predominantly in viral replication compartments (Figure 2).

**Figure 2.**
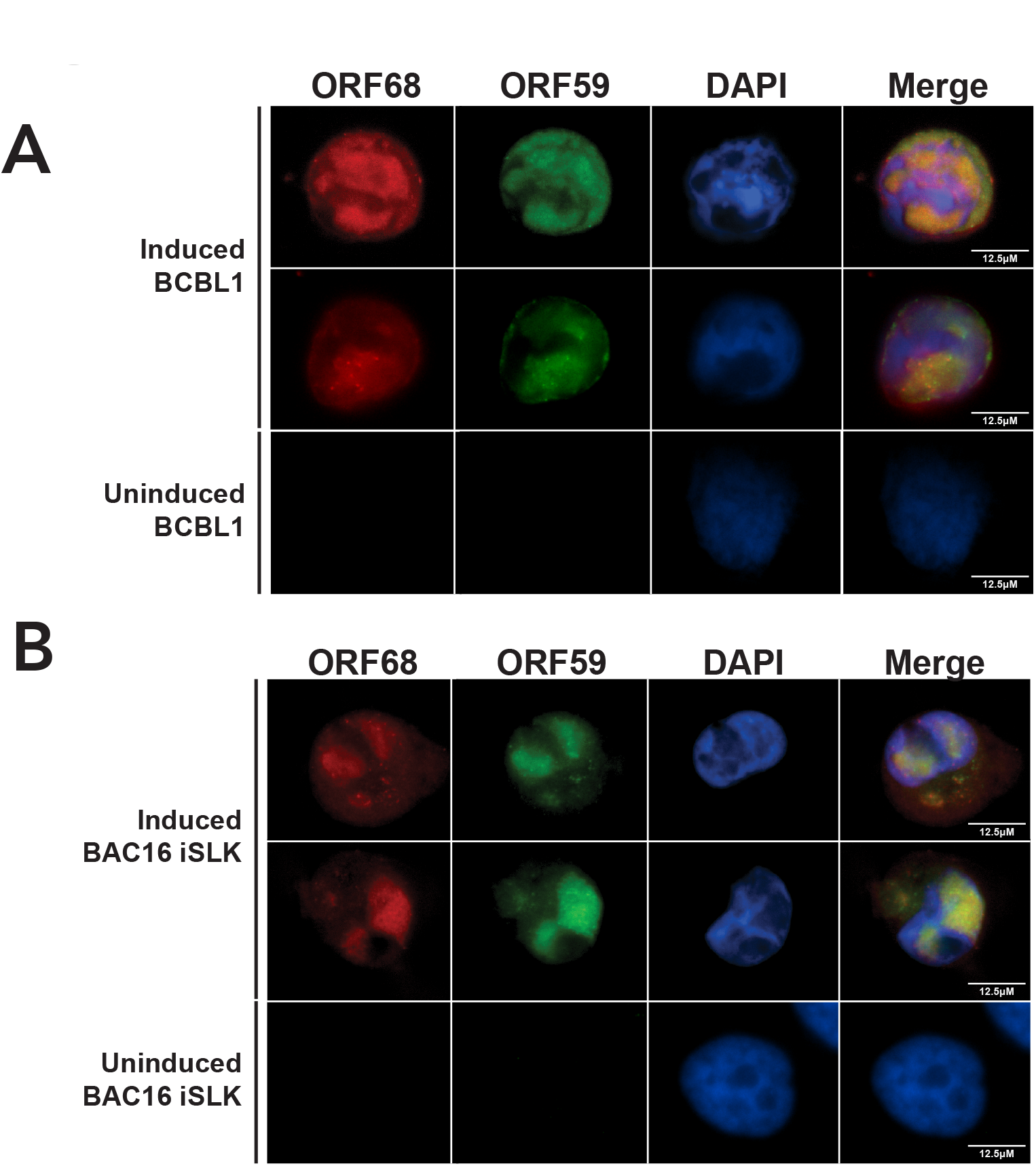
ORF68 localizes within replication compartments and in cytoplasmic puncta of KSHV infected cells. Reactivated and unreactivated BAC16.iSLK and TREx-BCBL1 cells were stained with antibodies to ORF68 (red), the viral replication compartment marker ORF59 (green), and DAPI (blue). The far-right panel shows the merged image.

### KSHV lacking ORF68 maintains viral DNA replication and late gene expression, but does not produce infectious virions

To evaluate the role of ORF68 in the viral replication cycle, we generated an ORF68-deficient virus (ORF68^PTC^) using the BAC16 Red Recombinase system (15) by inserting two premature termination codons 15 nt downstream of the translation start site. This virus was used to establish a latently-infected cell line in iSLK cells. To ensure that any observed phenotypes were not due to secondary mutations, we also engineered the corresponding mutant rescue virus (ORF68^PTC^-MR) (Figure 3a). Despite lacking ORF68 protein, the ORF68^PTC^ virus expressed both the delayed early protein ORF59 and the late protein K8.1 at levels similar to both the unmodified WT BAC16 KHSV and the ORF68^PTC^-MR virus (Figure 3b). Furthermore, qPCR analyses of viral DNA revealed no significant difference in DNA replication between any of the viruses (Figure 3c). We then monitored production of infectious progeny virions using a supernatant transfer assay, in which the supernatant from reactivated infected iSLK cells is filtered and transferred to target cells. Because the BAC16-derived KSHV contains a virally-encoded, constitutively-expressed GFP reporter gene, infected recipient cells can be quantified by GFP expression using flow cytometry. Here, ORF68^PTC^ showed a marked defect, as no virus production was detectable, while both WT and ORF68^PTC^-MR infected cells produced sufficient virus to infect nearly 100% of target cells (Figure 3d). Collectively, these data indicate that KSHV ORF68 is essential for a late stage event in the viral replication cycle.

**Figure 3.**
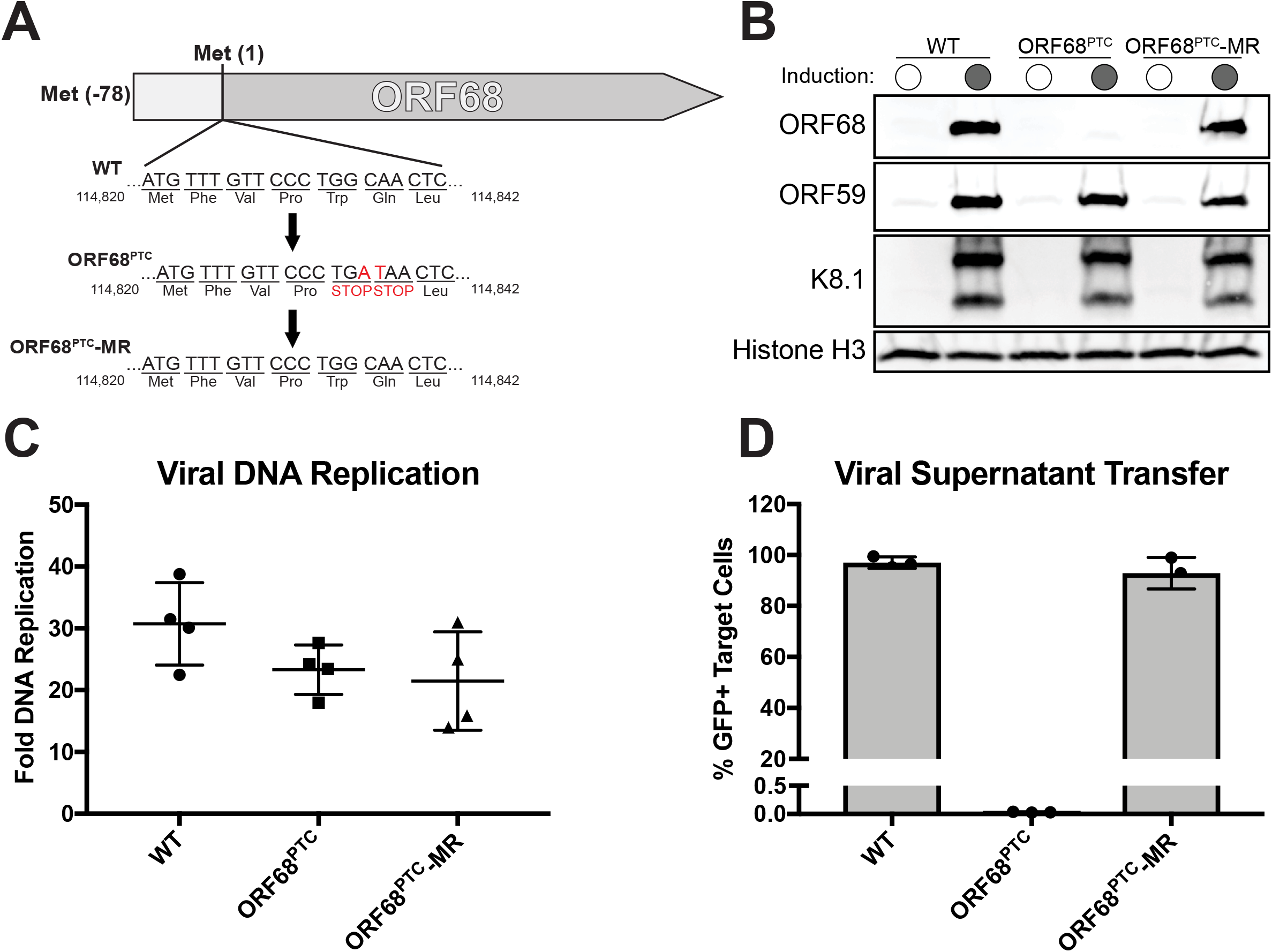
Characterization of the ORF68-deficient (ORF68^PTC^) mutant virus. (A) Diagram depicting the genetic locus of ORF68 and the mutations inserted into the ORF68^PTC^ virus. (B) Western blots showing expression of the early ORF59 protein and the late K8.1 protein in WT, ORF68^PTC^, and ORF68^PTC^-MR viruses. Histone H3 served as a loading control. (C) DNA replication was measured by qPCR of the viral genome before and after induction of the lytic cycle. (D) Progeny virion production was assayed by supernatant transfer and flow cytometry of target cells.

### In the absence of ORF68, KSHV DNA is not cleaved after replication

The above results, together with studies of alpha and betaherpesvirus ORF68 homologs (11, 14), suggested a role for KSHV ORF68 in viral DNA encapsidation. We therefore established an assay similar to what has been used in other herpesviruses to measure packaging-associated KSHV DNA cleavage, which occurs within the 801 bp GC-rich terminal repeat (TR) sequences present in 20 – 40 tandem copies in the KSHV genome (18) (Figure 4a). Briefly, DNA was isolated from infected iSLK cells and the TR sequences were released by digesting the DNA with Pstl-HF, which cleaves frequently within viral (and host) DNA, but not within the KSHV TRs. This should generate a ladder of TR sequences, representing the collection of individual cleavage events in the TRs that occur during packaging, which can be visualized by Southern blotting with a DIG-labeled TR probe. As expected, we observed a robust increase in the high MW uncleaved TRs following lytic reactivation of each sample, indicative of viral genome replication (Figure 4b). However, DNA from ORF68^PTC^ infected cells showed no evidence of TR cleavage, unlike DNA from WT and ORF68^PTC^-MR infected cells, which displayed the expected laddering phenotype (Figure 4b).

**Figure 4.**
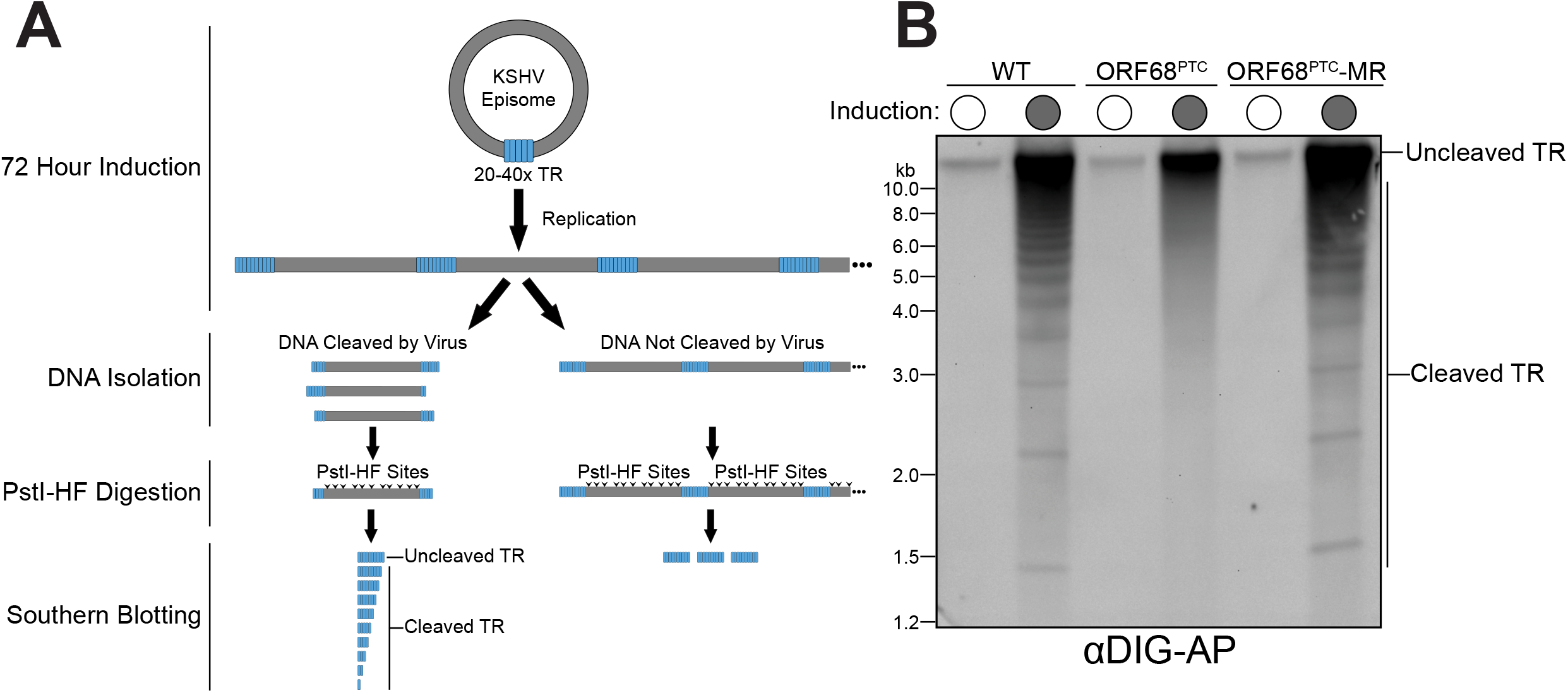
ORF68^PTC^ virus displays a genome cleavage defect. (A) Diagram depicting the DNA cleavage assay protocol and the expected TR DNA laddering phenotype (or lack thereof) upon infection with WT or ORF68^PTC^ virus. (B) Southern blot of the PstI-HF digested DNA with a TR probe from cells infected with WT, ORF68^PTC^, or ORF68^PTC^-MR virus.

### KSHV ORF68^PTC^ accumulates exclusively B-capsids

During herpesvirus assembly, immature procapsids are initially formed that contain internal scaffolding proteins, which are subsequently proteolytically cleaved and extruded during the process of DNA packaging (19, 20). This process yields mature, DNA-containing C-capsids, but in the case of unsuccessful DNA packaging, either empty A-capsids or scaffold-containing B-capsids are formed (3). To visualize the contribution of ORF68 towards capsid maturation, we performed transmission electron microscopy on fixed samples of lytically-reactivated iSLK cells containing either WT or ORF68^PTC^ KSHV. In the nuclei of WT KSHV infected cells, we observed A, B, and C capsids (Figure 5a, b). In contrast, ORF68^PTC^ infected cells contained exclusively B capsids (Figure 5a, b), indicating that KSHV DNA packaging is not initiated in the absence of ORF68.

**Figure 5.**
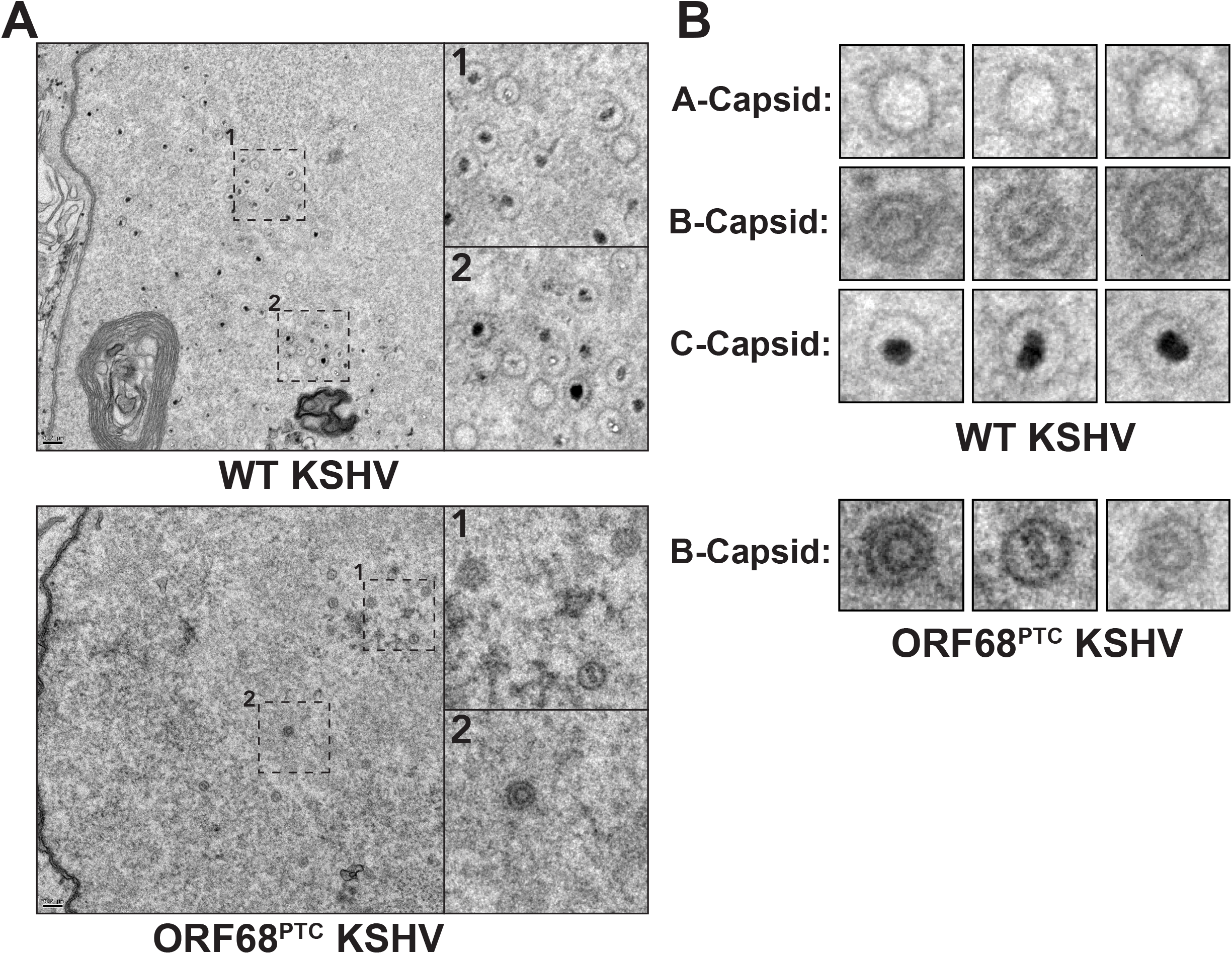
Transmission electron microscopy of WT and ORF68^PTC^ KSHV-infected cells. (A) Representative images of the nuclei of iSLK cells infected with WT or ORF68^PTC^ KSHV. Boxed regions are shown to the right at higher magnification to better visualize virion morphology. (B) Representative images of individual capsids observed in iSLK cells infected with WT or ORF68^PTC^ KSHV, grouped by their maturation stage. Only B-capsids were observed in cells infected with ORF68^PTC^ KSHV.

### ORF68 binds DNA with high affinity *in vitro* and is associated with metal-dependent nuclease activity

In order to study the biochemical activity of ORF68 as it relates to DNA packaging, we sought to generate pure ORF68 protein. Our attempts to isolate recombinant ORF68 from *E. coli* bacteria, *Pichia pastoris* yeast, and *SF9* insect cells did not yield properly folded protein (data not shown). Thus, we instead purified TwinStrep-tagged ORF68 from transiently transfected HEK293T cells by binding to Strep-TactinXT resin, followed by separation of ORF68 from the TwinStrep tag by PreScission protease cleavage and subsequent size exclusion chromatography (Figure 6a). Mass spectrometry of the purified sample detected ORF68 but no other contaminating proteins (data not shown). Notably, ORF68 eluted from gel filtration columns as a ^~^275 kDa oligomer (based on injected standards) rather than as a monomer (Figure 6b).

**Figure 6.**
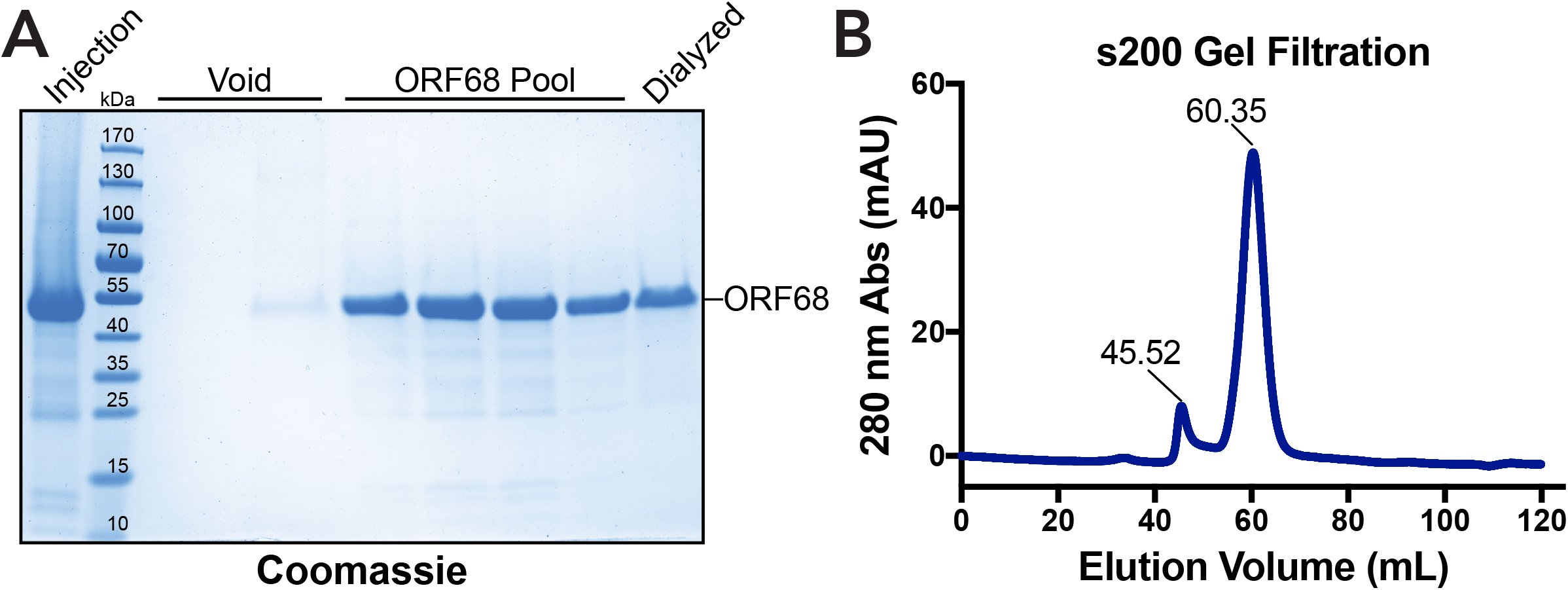
Purification of recombinant ORF68 from transiently transfected HEK293T cells. (A) Collodial coomassie-stained gel demonstrating the purity and molecular weight (^~^52 kDa) of ORF68 eluted from the s200 gel filtration column. Fractions containing ORF68 were pooled, concentrated, and dialyzed into storage buffer; a sample of which was loaded in the final lane. (B) Absorbance trace of the s200 gel filtration column, measured at 280 nm. The first peak represents the void elution at 45.52 ml and the second peak represents ORF68 at 60.35 ml.

Given the failed DNA cleavage and packaging phenotypes associated with the ORF68^PTC^ virus, we hypothesized that ORF68 might directly bind the viral genome to help direct DNA packaging. We therefore used electrophoretic mobility shift assays (EMSA) to monitor binding of purified ORF68 to a DNA probe containing the sequence of the KSHV TR *in vitro*. Indeed, ORF68 efficiently bound DNA with an apparent *K_d_* of (53.34 ± 2.11) nM (Figure 7a, b). The binding was cooperative, with a Hill coefficient of 1.88 ± 0.12 (Figure 7b). This is comparable to the 72 nM *K_d_* of KSHV LANA binding to the terminal repeat (21). Multiple higher-order binding events occurred at increased protein concentrations, resulting in several discrete bands of protein-bound DNA (Figure 7a). Although this *in vitro* DNA-binding is robust, it does not appear to be specific, as similar length probes of varying GC content from disparate sources were bound with similar affinity, and also displayed higher order binding events (data not shown).

**Figure 7.**
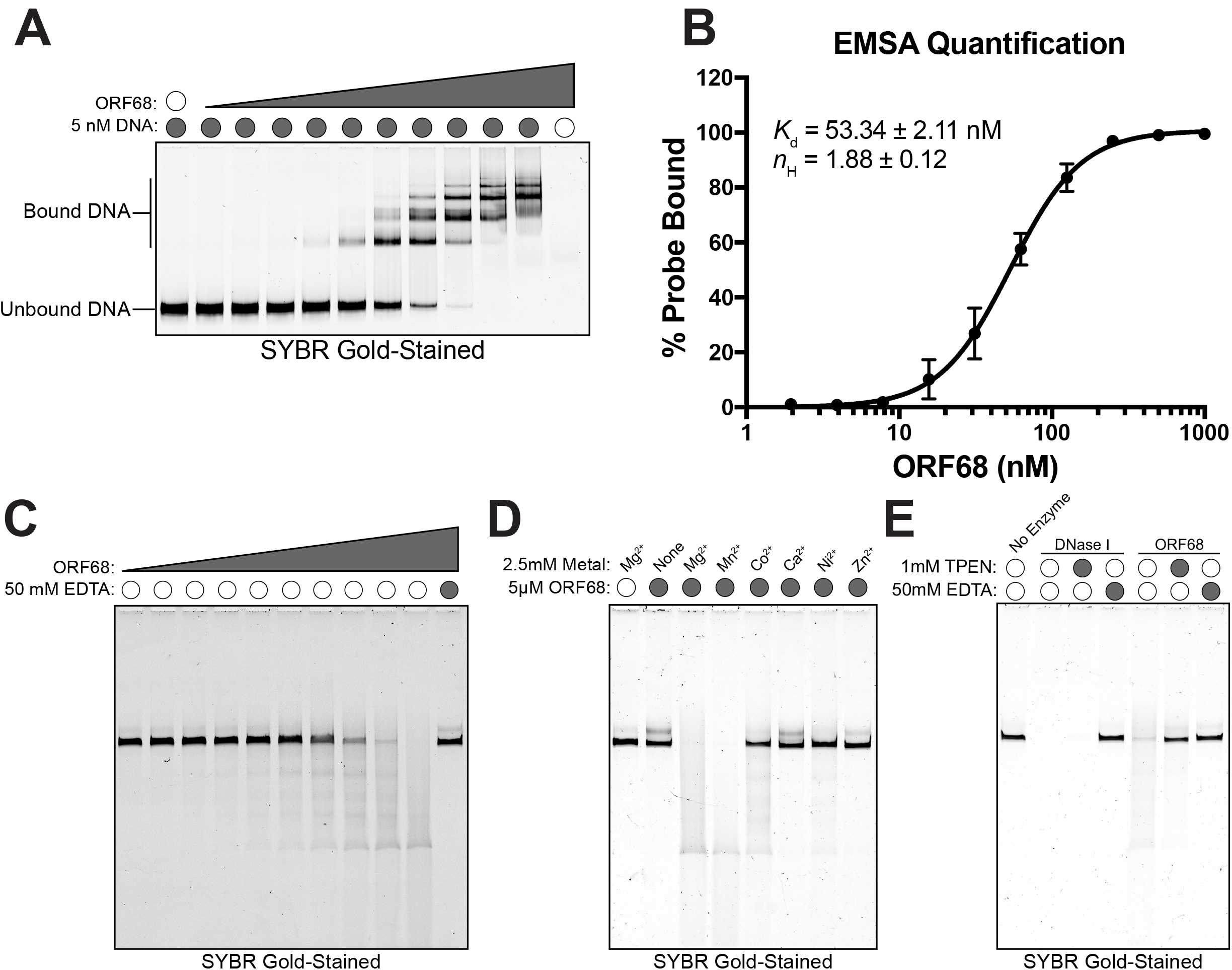
Purified recombinant ORF68 shows robust DNA binding and is associated with nuclease activity. (A) Representative EMSA showing increasing amounts of ORF68 binding to the subunit-length TR DNA probe. (B) Binding curve of ORF68 to DNA derived from quantification of three technical replicates of the EMSA, yielding an observed *K_d_* of (53.34 ± 2.11) nM with a Hill coefficient of 1.88 ±0.12. (C) Nuclease assay with increasing amounts of ORF68, incubated with an ^~^800bp TR probe in the presence or absence of EDTA. (D) Analysis of TR probe degradation in the presence of purified ORF68 upon addition of the indicated metal. (E) Gel comparing degradation of the TR probe in the presence of purified ORF68 or DNaseI +/− 1mM TPEN or 50 mM EDTA.

ORF68 has no identifiable domains aside from several predicted zinc finger motifs that appear to be important for the structural integrity of the protein (22). However, we noted that in the presence of Mg^2+^, the TR probe or other circular or linear plasmid DNA was degraded upon incubation with ORF68, suggesting that it may possess nuclease activity (Figure 7c, data not shown). Nuclease activity required metal ions for catalysis, with a preference for Mg^2+^ or Mn^2+^, drastically reduced activity with Co^2+^ or Ni^2+^, and no activity with Ca^2+^ or Zn^2+^ (Figure 7d). Reactions were sensitive to EDTA or similar metal-chelating agents, consistent with our observations that EDTA-containing buffers caused the precipitation of ORF68. Although our mass spectrometry analysis of purified ORF68 did not detect contaminating peptides, at present we cannot formally rule out the possibility that trace levels of a cellular nuclease co-purified with ORF68, rather than nuclease activity being an intrinsic feature of ORF68. However, given that ORF68 contains putative zinc finger motifs, we reasoned that in addition to inhibition by EDTA, the associated nuclease activity should be sensitive to the zinc-specific chelator N,N,N’,N’-tetrakis(2-pyridinylmethyl)-1,2-ethanediamine (TPEN), which has high affinity for Zn^2^+ (*K_d_*= 2.6 × 10^−16^ M), but very low affinity for Mg^2^+ (*K_d_*= 2.6 × 10^−2^ M) and other divalent cations (23–25). Indeed, the addition of TPEN prevented degradation of DNA in the presence of ORF68, but not DNase I, a nuclease that requires Mg^2+^ for catalysis but does not contain any structural zinc (Figure 7e). Collectively, these data demonstrate that KSHV ORF68 is a zinc finger-containing protein that can bind DNA and is associated with a zinc-dependent DNase activity.

## Discussion

Our data firmly establish that KSHV ORF68 is essential for genome encapsidation in infected cells, and further suggest novel roles for purified ORF68 protein in DNA binding and cleavage. This is the first demonstration of direct DNA binding by a DNA packaging-associated protein in gammaherpesviruses, and the ORF68-associated nuclease activity suggests a previously unappreciated function for this protein, which may relate to viral genome processing and encapsidation. Such a role would be consistent with the defects in viral TR cleavage, DNA packaging, and infectious virion formation we observed in cells infected with the ORF68^PTC^ virus. It is further supported by our TEM analysis of infected cells, which showed that in the absence of ORF68, DNA encapsidation was not initiated, as evidenced by the absence of A- and C-capsids. Finally, our data are in line with phenotypes observed upon deletion of the ORF68 homologs in HSV-1 (11), HCMV (14) and Epstein-Barr virus (EBV) (26). Indeed, in EBV, deletion of this BFLF1 gene was used to generate defective viral particles for use in vaccine development (26).

The ORF68 homolog in HSV-1 (UL32) was recently shown to be involved in modulation of disulfide bond formation during procapsid formation, which requires the presence of the conserved C-X-X-C motifs (11). Deletion of individual cysteines within four of the five motifs prevented complementation of a UL32 deletion mutant virus, and in the absence of UL32 the disulfide bond profiles of other viral proteins were altered. In ORF68, however, the C-X-X-C motifs appear to be important for its structural integrity, as chelation of Zn^2+^ causes ORF68 to precipitate during purification in a manner that suggests unfolding (unpublished observations). Thus, an important future challenge will be to resolve if these two functions are related, or if specific roles for UL32 and ORF68 have diverged between the alpha- and gammaherpesviruses.

In the KSHV episome, the terminal repeats are present as 20 to 40 copies of a highly repetitive ^~^800bp, 85% GC sequence (27). Unlike related herpesviruses, the packaging and cleavage signals in the KSHV terminal repeats are unknown. However, by using a probe generated from the entire repeat we could capture terminal repeats of any length. In a WT KSHV infection, this yields a ladder-like cleavage pattern reflective of the fact that the genome is replicated as head-to-tail concatemers, separated by an arbitrary number of terminal repeats. While each genome should be cut only once at each end during packaging, the specific number of repeats left at each genome terminus is variable (28, 29). This is in contrast to viruses lacking ORF68, which displayed WT levels of DNA replication by both qPCR and Southern blotting, but no apparent resolution of the amplified concatemers into unit-length genomes. These data reinforce the hypothesis that KSHV ORF68 is not required for genome replication but is integral to the process of DNA encapsidation. Interestingly, a recent cryo-electron microscopy (cryoEM) structure of the HSV-1 portal-vertex shows a pentameric assembly associated with the portal complex (30). Given that KSHV ORF68 purifies as an oligomer that is roughly the size of a pentamer, one possibility it that this cryoEM-visualized pentameric assembly may serve as an interaction site for ORF68 during the DNA packaging reaction.

Given the stage at which ORF68^PTC^ virus fails during virion maturation, we hypothesized that ORF68 may interact with the viral genome to promote DNA packaging. Indeed, purified ORF68 robustly bound the prototypical terminal repeat subunit, with an observed *K_d_* of ^~^50nM. This is comparable to the strength of the TR interaction for the KSHV latency-associated episome tethering protein LANA, strongly suggesting that viral DNA binding is a biologically relevant feature of ORF68 (21). However, it is important to note that *in vitro*, the DNA binding activity of ORF68 does not have apparent sequence specificity. In cells, it is possible that sequence specificity is conferred by other viral (or cellular) factors. It is also possible that the DNA binding activity participates in a distinct facet of the viral lifecycle, although we did not detect any defects in the ORF68^PTC^ virus prior to encapsidation. The ^~^800bp terminal repeat subunits are somewhat long for traditional EMSA probes and as such, multiple binding events could be observed. Furthermore, binding of ORF68 to DNA was cooperative, as is frequently observed for DNA binding proteins (31). This cooperativity may indicate that ORF68 binds the DNA probe at multiple sites, that the binding of the first ORF68 molecule alters the DNA conformation in a way that promotes additional ORF68 binding, or simply reflect our observation that ORF68 exists as an oligomer in solution.

In addition to DNA binding, we made the unexpected observation that in the presence of the divalent cations Mg^2+^ or Mn^2+^, incubation with ORF68 caused degradation of DNA. These cations are commonly used by nucleases (32), and future studies are aimed at exploring residues in ORF68 that could serve to coordinate divalent cations. Our robust purification scheme, coupled with the absence of any detectable co-purifying peptides by mass spectrometry and the fact that addition of the zinc-chelator TPEN specifically blocked ORF68-associated nuclease activity suggest that the activity is intrinsic to ORF68. However, aside from the zinc finger motifs, no predicted domains or structural information is currently available for ORF68. Thus, a key challenge will be to establish if and how ORF68 promotes DNA cleavage in cells. Formal proof of nuclease activity will ultimately require structural data to identify putative catalytic domain(s), which should enable isolation of point mutants to be tested for DNA cleavage. Similar to the DNA binding activity, our *in vitro* cleavage reactions do not capture TR-specific targeting, as we found that other non-viral DNA was similarly susceptible to ORF68-associated degradation (unpublished observations). This might suggest that other components of the packaging machinery are required to confer specificity or, alternatively, that the putative nuclease activity is involved in an ORF68 function independent of DNA encapsidation.

While seven herpesviral proteins are required for proper DNA encapsidation, only two have been shown to directly act on DNA. A crystal structure of a fragment of the HSV-1 UL15 terminase motor protein, which is homologous to KSHV ORF29, revealed that it adopts an RNase H-like fold and possess non-specific DNA nicking *in vitro* (8, 33). Purified HSV-1 UL28, another terminase motor protein and the homolog of KSHV ORF7, has also been shown to bind terminal repeat DNA *in vitro* (7). It has high specificity for certain terminal repeat DNA probes, but only after heat treatment of the DNA, which causes them to adopt non-duplex structures (7). This suggests that the UL28-DNA interaction is structure dependent, in contrast to KSHV ORF68, which bound duplex DNA with no apparent sequence specificity or requirement for higher order structures. Whether the UL15 and UL28 functions are conserved in their KSHV homologs ORF29 and ORF7, and how KSHV ORF68 mechanistically contributes to DNA encapsidation remain to be established.

Finally, although the majority of ORF68 is concentrated in replication compartments, the protein also localizes within discrete puncta in the cytoplasm, as well as in larger aggregates adjacent to the nucleus. While we hypothesize that the larger structures represent unfolded protein in aggresomes, the smaller punctate structures are suggestive of a distinct function for ORF68 in the cytoplasm, possibly unrelated to its role in packaging. Notably, HSV-1 UL32 was also shown to localize to the cytoplasm, particularly late in infection, where it forms perinuclear foci (11). Given the frequently multifunctional roles of viral proteins, it will be of interest to explore possible additional cytoplasmic activities of ORF68.

## Methods

### Plasmids

TwinStrep-PreScission-ORF68 was cloned into the pHEK293 UltraExpression I vector (Clontech) digested with XhoI and Sall. ORF68 was amplified with the primers 5’-TGGAATTCTGCAGATATGTTTGTTCCCTGGCAACTCG-3’ and 5’-GCCACTGTGCTGGATTCAAGCGTACAAGTGTGACGTCT-3’ while TwinStrep-PreScission was amplified using the primers 5’-CCTCCCCGGGCTCGAATGAGTGCGTGGAGTCATCCTCAATTCGAGAAAGGTG-3’ and 5’-GGGCCCCTGGAACAGAACTTCCAGTCCGGATTTTTCGAACTGCGGG-3’. The two fragments were first assembled by PCR and then inserted into the vector by InFusion cloning (Clontech).

ORF68 and ORF68-Extended were cloned into pCDNA4 (ThermoFisher) digested with EcoRV using 5’-TGGAATTCTGCAGATATGTTTGTTCCCTGGCAACTCG-3’ for ORF68, 5’-TGGAATTCTGCAGATATGTCACGAGGCAGAAGCTGG-3’ for ORF68-Extended, and 5’-GCCACTGTGCTGGATTCAAGCGTACAAGTGTGACGTCT-3’ as the reverse primer for both. PCR products were inserted into the vector using InFusion cloning.

### Cell Lines

HEK293T (ATCC CRL-3216) cells were maintained in DMEM +10% FBS. HEK293T-ORF68 cells were maintained in DMEM + 10% FBS, 325 μg/ml zeocin. iSLK.BAC16 (15) cells were maintained in DMEM + 10% FBS, 1 mg/ml hygromycin B, and 1 μg/ml puromycin. TREx-BCBL1 (34) cells were maintained in RPMI 1640 + 20% FBS.

### Viral mutagenesis and infection studies

The KSHV ORF68 Premature Termination Codon (ORF68^PTC^) mutant and corresponding mutant rescue (MR) were engineered using the scarless Red recombination system in BAC16 GS1783 *E. coli* as previously described (15), except using two gBlocks (IDT) to introduce the mutation. Each gBlock contained half of the kanamycin resistance cassette, as well as the desired mutation, and were joined by short overlap extension PCR before being used as the linear insert in the established protocol.

The BAC16 ORF68 mutant and MR were purified using the NucleoBond BAC 100 kit (Clontech). iSLK cell lines latently infected with the KSHV ORF68^PTC^ virus were then established by co-culture of the relevant BAC16-containing HEK293T cells with uninfected target iSLK-Puro cells. HEK293T cells constitutively expressing ORF68 (HEK293T-ORF68) were transfected with 12 μg of BAC16 containing ORF68^PTC^ or MR using linear polyethylenimine (PEI, MW ^~^25,000) at a 1:3 DNA:PEI ratio. The following day, the cells were trypsinized and mixed 1:1 with iSLK-puro cells, then treated with 30 nM 12-O-Tetradecanoylphorbol-13-acetate (TPA) and 300 nM sodium butyrate for 4 days to induce lytic replication. Cells were then incubated with selection media containing 300 μg/ml hygromycin B, 1 μg/ml puromycin, and 250 μg/ml G418. Media was replaced every other day for ^~^2 weeks, gradually increasing the hygromycin B concentration to 1 mg/ml until there were no HEK293T cells remaining and the iSLK cells were green and replicating.

For induction studies, BAC16-containing iSLK cells were treated with 1 μg/ml doxycycline and 1 mM sodium butyrate, and TREx-BCBL1 cells were induced with doxycycline (1 μg/ml) and TPA (25 ng/ml) for the indicated amount of time. Virion production from reactivated iSLK cells was measured using supernatant transfer assays at 72 hours post induction. The supernatant was filtered through a 0.45 μM PES filter, then 2 ml were mixed with 1×10^6^ freshly trypsinized HEK293T cells. The cell mixture was placed into a 6 well plate and centrifuged at 1,200 × *g* for 2 hours at 32°C. The following day, cells were trypsinized, fixed in 4% paraformaldehyde, then analyzed for GFP expression by flow cytometry (BD Csampler, BD Biosciences). Data was analyzed using FlowJo v10.

### Generation of HEK293T cells stably expressing ORF68

HEK293T-ORF68 cells were generated by cloning ORF68 into pLJM1 (a gift from David Sabatini (Addgene plasmid #19319) (35)) in which the puromycin resistance had been exchanged for zeocin. The vector was digested with AgeI and EcoRI (New England Biolabs) and ORF68 was amplified by PCR using primers 5’-CGCTAGCGCTACCGGATGTTTGTTCCCTGGCAACTCGG-3’ and 5’-TCGAGGTCGAGAATTTCAAGCGTACAAGTGTGACGTCTG-3’ and inserted using T4 DNA Ligase (NEB). 1.6 μg of this plasmid was co-transfected into HEK293T cells along with the lentiviral packaging plasmids pMD2.G (0.5 μg) and psPAX2 (1.3 μg) (gifts from Didier Trono (Addgene plasmid #12259 and #12260)) using PEI. 48 hours after transfection, the supernatants were collected, filtered through a 0.45 μm PES filter, diluted 1:4 (supernatant:media) in serum-free DMEM containing 8 μg/ml of polybrene, and added to the wells of a 6 well plate containing 1×10^6^ HEK293T cells per well. The plates were centrifuged for 2 hours at 1,200 × *g* (32°C), incubated overnight, and the media replaced with fresh DMEM (10% FBS) containing 325 μg/ml zeocin until no further cell death was observed, roughly two weeks.

### Western blots and antibody production

Cells were lysed in protein lysis buffer (50 mM Tris-HCl pH 7.6, 150 mM NaCl, 3 mM MgCl_2_, 10% glycerol, 0.5% NP-40, cOmplete EDTA-free Protease Inhibitors [Roche]) and clarified by centrifugation at 20,000 × *g* for 10 min at 4°C. Lysates were resolved by SDS-PAGE and western blotted with rabbit anti-ORF68 (1:5000), rabbit anti-ORF59 (1:10,000), rabbit anti-K8.1 (1:10,000), rabbit anti-histone H3 (1:3000, Cell Signaling 4499S), or mouse anti-GAPDH (1:5000, Abcam ab8245). Rabbit anti-ORF68, -ORF59, and -K8.1 was produced by Pocono Rabbit Farm & Laboratory by immunizing rabbits against MBP-ORF68, -ORF59, or -K8.1 (gifts of Denise Whitby, (36)). Sera was harvested and used directly for ORF59 and K8.1, while ORF68-specific antibodies were isolated by first collecting antibodies that bound an MBP-ORF68 column, followed by removal of non-specific antibodies on an MBP column. Both selection columns were generated using the AminoLink Plus Immobilization Kit (ThermoFisher).

### Immunofluorescence assays

Cells were fixed with 4% paraformaldehyde, permeabilized with ice-cold methanol at -20°C for 20 min, then blocked with BSA blocking buffer (PBS, 1% Triton X-100, 0.5% Tween20, 3% BSA) for 30 minutes at room temperature. They were then incubated with rabbit anti-ORF68 and mouse anti-ORF59 (Advanced Biotechnologies) antibodies (diluted 1:100 in blocking buffer) overnight at 4°C, washed with PBS, and incubated with secondary antibody (AlexaFluor594 or DyLight650, 1:1000 in BSA blocking buffer) for 1 h at 37°C, and mounted using VectaShield HardSet with DAPI (ThermoFisher). Images were acquired on an EVOS FL inverted fluorescent microscope (ThermoFisher).

### DNA isolation and qPCR

iSLK-BAC16 cells were incubated with 5x proteinase K digestion buffer (50 mM Tris-HCl pH 7.4, 500 mM NaCl, 5 mM EDTA, 2.5% SDS) and digested with proteinase K (80 μg/ml) overnight at 55°C. The gDNA was isolated using Zymo Quick gDNA Miniprep Kit according to the manufacturer’s instructions.

Quantitative PCR was performed on the isolated DNA using iTaq Universal SYBR Green Supermix on a QuantStudio3 Real-Time PCR machine. DNA levels were quantified using relative standard curves with primers specific for KSHV ORF57 (5’-GGTGTGTCTGACGCCGTAAAG-3’ and 5’-CCTGTCCGTAAACACCTCCG-3’) or a region in the GAPDH promoter (5’-TACTAGCGGTTTTACGGGCG-3’ and 5’-TCGAACAGGAGGAGCAGAGAGCGA-3’). The relative genome numbers were normalized to GAPDH to account for loading differences and to uninduced samples to account for starting genome copy number.

### Protein purification

TwinStrep-PreScission-ORF68 was transfected using PEI into 50 90% confluent 15 cm plates of HEK293T cells for 48h. Cells were lysed in 5 ml of lysis buffer (100 mM Tris-HCl pH 8.0, 300 mM NaCl, 1 mM DTT, 5% glycerol, 0.1% CHAPS, 1 μg/ml avidin, cOmplete EDTA-free protease inhibitors [Sigma-Aldrich]) per 1 g of cell pellet and rotated at 4°C for an additional 30 minutes. The viscosity was reduced by sonication, whereupon debris was removed by centrifugation at 20,000 × *g* (4°C) for 30 min followed by filtration through a 0.45 μm PES filter with a glass pre-filter (Millipore). Lysate was passed three times over a gravity column containing Strep-Tactin XT Superflow slurry (IBA) then washed with 5 column volumes of strep running buffer (100 mM Tris-HCl pH 8.0, 300 mM NaCl, 1 mM DTT, 5% glycerol, 0.1% CHAPS). ORF68 was eluted by rotating the column overnight in HRV 3C Protease (Millipore) at a 1:100 Protease:ORF68 ratio in buffer containing 2 mM DTT, then concentrated to ^~^500 μL using a 30 kDa cut-off Centriprep spin concentrator (Millipore). The concentrated eluate was centrifuged at 20,000 × *g* for 10 min to remove precipitates and injected onto a HiLoad 16/600 Superdex 200 pg gel filtration column (GE Healthcare), previously equilibrated with Gel Filtration buffer (50 mM HEPES pH 7.6, 100 mM NaCl, 100 mM KCl, 5% glycerol, 0.1% CHAPS). Gel filtration was performed using an AKTA Pure FPLC (GE Healthcare). Fractions resulting from the gel filtration were analyzed by SDS-PAGE and staining with colloidal Coomassie (37). Fractions containing ORF68 were pooled and concentrated as before, until the total concentration was ^~^2 mg/mL as evaluated by absorbance at 280 nm. The concentrated protein was injected into a 20 kDa cut-off Slide-A-Lyzer dialysis cassette (ThermoFisher) and dialyzed against storage buffer (Gel Filtration buffer lacking CHAPS). Dialyzed protein was diluted to 1 mg/ml, aliquoted, and snap-frozen in liquid nitrogen before being stored at −80°C.

### Electrophoretic mobility shift assay

Single-subunit terminal repeat DNA probes were generated by digesting pK8TR (a gift of the Kaye lab (18)) with AscI overnight. The resulting cleavage products were separated on an agarose gel and the 800bp band was excised, isolated from agarose, and purified further by phenol:chloroform extraction and ethanol precipitation.

EMSA assays were assembled on ice in gel shift buffer (25 mM HEPES pH 7.6, 50 mM NaCl, 50 mM KCl, 0.1 mg/ml BSA, 0.05% CHAPS, 10% glycerol) with 5 nM probe and 2-fold dilutions of ORF68 ranging from 1.95 nM to 1000 nM. Once assembled, the reactions were incubated at 30°C for 30 minutes. Samples were directly loaded onto a 3.5% 29:1 polyacrylamide:bisacrylamide native gel containing 45 mM Tris-borate pH 8.3, 10% glycerol, which was degassed thoroughly before polymerization. Samples were separated by electrophoresis at 250V for 1 hour at 4°C in pre-chilled 0.5X TB (45 mM Tris-borate pH 8.3). Tris-borate buffer was used in lieu of TBE because ORF68 precipitates in the presence of EDTA. The native polyacrylamide gel was cast to a final concentration of 0.5X Tris-borate, 10% glycerol, and was degassed thoroughly before polymerization. Following the electrophoresis, the gel was removed from the running cassette and stained with SYBRgold (ThermoFisher) in 0.5X TB for 30 minutes, rocking at room temperature and protected from light. Staining solution was removed by briefly washing with fresh 0.5X TB and the gel was imaged on a ChemiDoc Touch (Bio-rad). All bands were quantified using ImageLab v6.0 (Bio-rad) and percent bound probe was determined by dividing shifted band intensity by the total intensity of all bands in that lane. Binding curves were generated using Prism v7 (GraphPad) by non-linear regression fit using least-squares. Shown data represents three technical replicates performed on different days.

### Nuclease assays

The EMSA probe was also used for nuclease assays and was produced as described above. Reactions were assembled in nuclease buffer (25 mM HEPES pH 7.6, 50 mM NaCl, 50 mM KCl, 0.1 mg/ml BSA, 2.5 mM MgCl_2_) with 1 nM DNA probe and 2-fold dilutions of ORF68 ranging from 9.77 to 5000 nM. The reactions were incubated at 37°C overnight. The following day, reactions were quenched using stop buffer (4 M urea, 50 mM EDTA, 1 mM CaCl_2_, 0.8 U proteinase K [New England Biolabs], 1x Purple Loading dye [New England Biolabs]; final concentrations) and incubating at room temperature for 30 minutes. The samples were then loaded onto a denaturing polyacrylamide gel (8 M Urea, 4.5% 29:1 acrylamide:bisacrylamide, 90 mM Tris-borate pH 8.3, 2 mM EDTA, degassed before polymerization), which had been pre-run at 200V for 30 minutes in 1X TBE to evenly heat the gel. The samples were separated at 200V for 45 minutes before being removed from the running cassette and stained with SYBRgold in 1X TBE for 30 minutes at room temperature, protected from light. Gels were visualized using a ChemiDoc Touch (Bio-Rad). Metal-dependence tests were performed as described but changing the 2.5 mM MgCl_2_ to the chloride salts of the indicated divalent cations. TPEN-treatment was performed as described but protein was treated with either TPEN in 100% ethanol (1 mM), mock treated with 100% ethanol, or EDTA (50 mM) for 10 minutes on ice prior to adding the DNA substrate. DNaseI (New England Biolabs) was diluted to 0.02U in ORF68 storage buffer before addition to reactions.

### Southern blotting

BAC16.iSLK cells were harvested in proteinase K digestion buffer and incubated overnight with proteinase K (80 μg/ml). DNA was isolated by phenol:chloroform extraction and ethanol precipitation, then resuspended in TE buffer. DNA (10 μg) was digested with PstI-HF (New England Biolabs) overnight then separated by electrophoresis in a 0.7% agarose 1x TBE gel stained with SYBRsafe. The DNA was transferred to a NYTRAN-N membrane (GE Healthcare) by capillary action in 20x SSC overnight and crosslinked to the membrane in a StrataLinker 2400 (Stratagene) using the AutoUV setting.

The membrane was treated according to the DIG High Prime DNA Labeling and Detection Starter Kit II (Roche) following the manufacturer’s instructions. Briefly, the membrane was hybridized at 42°C in EasyHyb Buffer with a DIG-labeled linear terminal repeat subunit derived from AscI (New England Biolabs) digested pK8TR, which was labeled according to kit instructions. Following overnight hybridization, the membrane was washed twice with 2X SSC (+0.1% SDS) at room temperature and then twice with 0.5X SSC (+0.1% SDS) at 68°C. Using supplied reagents, the blot was then blocked, incubated with anti-DIG-AP antibody, and visualized on a ChemiDoc Touch (Bio-Rad).

### Electron microscopy

iSLK.BAC16 WT or ORF68^PTC^-infected cells were induced for 48 hours as described above. Cells were washed once with PBS and fixed with 2% glutaraldehyde, 4%paraformaldehyde in 100 mM sodium cacodylate buffer, pH 7.2 for 10 minutes at room temperature. Fixative was removed and cells were scraped into 2% glutaraldehyde in 100 mM sodium cacodylate buffer, pH 7.2 and evenly resuspended by pipetting. Cells were then washed with 1% osmium tetroxide, 0.8% ferricyanide in 100 mM sodium cacodylate buffer, pH 7.2, before being treated with 1% uranyl acetate and dehydrated with increasing concentrations of acetone. Samples were then infiltrated and embedded in resin, from which 70 nm sections were cut on a Reichert-Jung microtome. Sections were picked up on copper mesh grids coated with 0.5% formvar, post-stained with uranyl acetate and lead citrate, and examined using an FEI Tecnai 12 transmission electron microscope.

## Acknowledgements

We thank all members of the Glaunsinger and Coscoy labs, in particular Angelica Castañeda and Allison Didychuk, as well as Dr. Sandra Weller for their helpful suggestions and critical reading of the manuscript. We also thank Reena Zalpuri and all of the staff at the University of California Berkeley Electron Microscope Laboratory for advice and assistance in electron microscopy sample preparation and data collection. B.G. is an investigator of the Howard Hughes Medical Institute. This research was also supported by NIH R01AI122528 to B.G.

